# Multidimensional dynamics of the proteome in the neurodegenerative and ageing mammalian brain

**DOI:** 10.1101/472118

**Authors:** Byron Andrews, Sarah Maslen, Leonardo Almeida-Souza, J Mark Skehel, René Frank

**Affiliations:** MRC Laboratory of Molecular Biology, Francis Crick Avenue, Cambridge, CB2 0QH, United Kingdom; Storm Therapeutics, Babraham Research Campus, Cambridge CB22 3AT, United Kingdom; School of Biomedical Sciences, Faculty of Biological Sciences, University of Leeds, LS2 9JT

## Abstract

The amount of any given protein in the brain is determined by the rates of its synthesis and destruction, which are regulated by different cellular mechanisms. Here, we combine metabolic labelling in live mice with global proteomic profiling to simultaneously quantify both the flux and amount of proteins in mouse models of neurodegeneration. In multiple models, protein turnover increases were associated with increasing pathology. This method distinguishes changes in protein expression mediated by synthesis from those mediated by degradation. In the *App^NL-F^* knockin mouse model of Alzheimer’s disease increased turnover resulted from imbalances in both synthesis and degradation, converging on proteins associated with synaptic vesicle recycling (Dnm1, Cltc, Rims1) and mitochondria (Fis1, Ndufv1). In contrast to disease models, ageing in wildtype mice caused a widespread decrease in protein recycling associated with a decrease in autophagic flux. This simple multidimensional approach enables the comprehensive mapping of proteome dynamics and identifies affected proteins in mouse models of disease and other live animal test settings.

## INTRODUCTION

In the mammalian brain, the rate of protein turnover ranges from minutes to several days (Dwyer et al., 1980; Price et al., 2010b; Savas et al., 2012; Toyama et al., 2013). This extraordinary flux poses a particular challenge for the brain because information must outlive the molecular substrates in which they are stored (Crick, 1984; Lisman et al., 2012). In the adult brain almost all neurons are post-mitotic, thus turnover and repair mechanisms are not helped by replication during cell division, as applies in many other tissues. The rate of proteome turnover is regulated by multiple factors and mechanisms, including ubiquitin-proteasome and autophagy-mediated degradation (Goldberg, 2003; Ohsumi, 2006; Vilchez et al., 2014). Protein turnover perturbations cause severe neurological dysfunction (Harris and Rubinsztein, 2011). Indeed, the most common neurodegenerative diseases are characterized by imbalances in the turnover of a few proteins, resulting in their accumulation into mis-folded protein aggregates (Eisenberg and Jucker, 2012; Goedert, 2015). These inclusions appear to be resistant to cellular mechanisms of repair (Ye et al., 2015). Abnormal inclusions in non-neuronal cell culture have been shown to have widespread impact on the proteome and its functions (Kim et al., 2016; Olzscha et al., 2011; Woerner et al., 2016). However, it is not known if neurodegenerative diseases have an impact on global proteome turnover in the mammalian brain.

Neurodegenerative diseases are characterized by synapse loss, cognitive decline, and eventual neuronal death. What triggers these diseases is unknown, except for a very small subset that are caused by familial mutations (Guerreiro and Hardy, 2014; Yancopoulou and Spillantini, 2003). Yet, even in these rare cases, a comprehensive understanding of what downstream pathological pathways are involved in cognitive decline, synapse and neuronal loss is lacking (De Strooper and Karran, 2016).

In most neurodegenerative diseases, including Alzheimer’s disease (AD), it is apparent that pathology arises over many years, perhaps decades (H. Braak and E. Braak, 1991; Dubois et al., 2016; Goedert, 2015). Consequently, pathology in the early stages of the disease could be masked by increased repair and adaptation (Hardy et al., 2014; Masters and Selkoe, 2012; Musiek and Holtzman, 2015; Zetterberg and Mattsson, 2014). Therefore, methods capable of detecting these changes in repair could indicate the earliest upstream pathways of the disease (De Strooper and Karran, 2016).

To study AD, mouse models provide an excellent setting because genetic approaches can be applied within an organism that is neuroanatomically and molecularly similar to humans (Bayés et al., 2011). Many useful models are available, that show varying signs of cognitive decline and synaptic loss, but none reflect the full cascade of pathology including neuronal death (Jucker, 2010). Thus, approaches are required that can reconcile the range of molecular abnormalities in different mouse models of disease and identify affected molecular pathways.

*In vivo* metabolic labelling and global proteomic profiling has the capacity to measure the dynamics of individual proteins throughout the proteome (Hinkson and Elias, 2011; Krüger et al., 2008; Larance and Lamond, 2015; McClatchy et al., 2007; Schwanhäusser et al., 2011). Here, we first established a method using ^13^C heavy lysine (K6) labelling to detect global proteome turnover change in mice. Next, we developed the assay to simultaneously measure changes in protein turnover and expression level *in vivo*. This multiplex screen of proteome dynamics is applicable to any protein in any tissue and distinguishes between changes driven by synthesis or degradation of a protein. We applied this screen to quantify ~1000 proteins in three mouse models of neurodegenerative disease at pre-symptomatic and symptomatic ages. In all models that we tested, increased neuropathology is associated with increased protein turnover and changes in the amount of some specific proteins, caused by measurable alterations in their synthesis or degradation. Finally, we used the method to investigate the proteome dynamics that are associated with ageing in healthy mice. Global protein turnover decreased with age, which was associated with a slowdown in autophagy. This resource reveals novel signatures of pathology, facilitates comparisons between different mouse models of disease and contrasts neurodegeneration with the mechanisms of ageing.

## RESULTS

### Detecting proteome turnover in mouse models of disease

There are multiple approaches capable of measuring protein turnover (Doherty et al., 2005; Heo et al., 2018; Larance et al., 2011; Price et al., 2010a; Savitski et al., 2018). We devised a simple approach that matched the following three criteria: 1) Requires minimal experimental design, 2) is amenable to cohorts of multiple test and control mice, and 3) enables straightforward identification of peptides, label incorporation and turnover. To quantify changes in protein turnover, mice were fed a diet in which the essential amino acid, lysine (K0), was replaced with a ^13^C stable isotope derivative (K6) for 6-8 days (Figure 1A). The rate of K6 incorporation was directly quantified by the ratio of K6 to K0 in each mouse. We benchmarked the accuracy of identifying heavy-light peptide pairs using a decoy unlabelled dataset, which indicated an empirical FDR of 2% and 0.01% using single and double counting of K0-K6 pairs, respectively. Therefore, double counts were used throughout.

**Figure 1.**
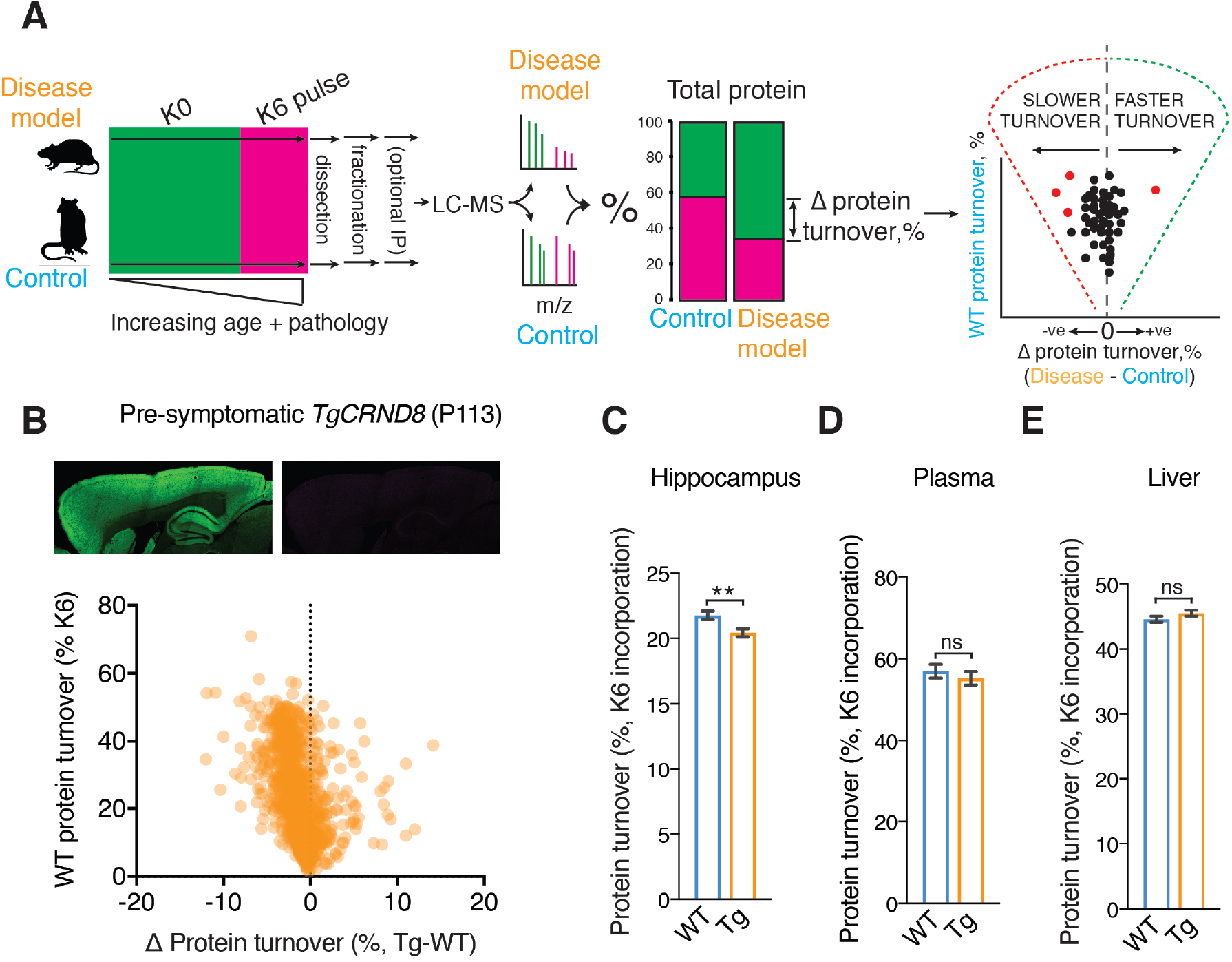
Metabolic labelling of live mice to measure changes in protein turnover. **A, *left*,** Schematic summarising the ^13^C heavy lysine (K6) labelling method for measuring changes in protein turnover in mice. Cohorts of genetically and age matched mice were maintained on regular food until the desired labelling window. The groups of mice are then switched to K6 diet for an identical period before tissues were processed for orbitrap LC-MS/MS. ***Middle***, Ion pairs with a 6 Da difference in mass were detected and sequenced, corresponding to peptides with and without K6. The relative incorporation of K6 was calculated in disease and wildtype mice. ***Right***, The mean difference in K6 incorporation is calculated for each protein and plotted (*x*-axis) to highlight slowdown or increases in protein turnover versus whether a protein has fast or slow turnover (WT K6 incorporation, *y*-axis). **B, *Top*,** Immunohistochemical detection of synaptic marker, left, Psd95 and right, β-amyloid pathology in sagittal sections of pre-symptomatic *TgCRND8* (P113) mouse brain. ***Bottom***, Scatter plot showing protein turnover changes in the hippocampus of pre-symptomatic (P113) *TgCRND8* mice (see Table S1). 1392 proteins were quantified in both diseased and healthy cohorts of mice (2 of 3 mice). The mean difference of K6 incorporation for each protein (*x*-axis, Tg - WT) was plotted against the incorporation in WT (*y*-axis). 3 *TgCRND8* and 3 age-, sex-, and background matched WT mice were K6 labelled for 6 days. These data indicate a global decrease in protein turnover for hippocampal proteins with both a slow and fast turnover. **C** Bar chart showing average protein turnover of 1392 proteins in the hippocampus of *TgCRND8* and WT mice. An overall 6.1% slowdown in turnover was recorded in the hippocampus (*P* = 0.0037). Error bars indicate SEM. ** *P* < 0.01. **D**, Bar chart showing average plasma protein turnover in *TgCRND8* and WT mice. No significant difference was detected (*P* = 0.128, n = 118). Error bars indicate SEM. ns, not significant. **E**, Bar chart showing average liver protein turnover in *TgCRND8* and WT mice. No significant difference was detected (*P* = 0.17, n = 590). ns, not significant.

To validate this simple method of measuring changes in protein turnover and its applicability to neurodegenerative disease we used *TgCRND8* mice, an aggressive transgenic mouse model of familial Alzheimer’s disease (AD) that overexpresses hAPP (Chishti et al., 2001). This mouse line develops several pathologies characteristic of AD including β-amyloid plaques, synapse loss, and behavioural phenotypes (Chishti et al., 2001). Three 3-month-old *TgCRND8* and three age-, sex- and genetic background-matched control mice were labelled with K6 food for 6 days. At the end of this labelling period the hippocampi from these mice were collected for LC-MS (Figure 1A). Measuring the change in K6 incorporation between disease and control tissues gave a snapshot of proteome turnover associated with the disease.

In each tissue sample from each mouse we identified an average of 72,261 peptides (±4,231 sd), with a total of 130,516 unique peptides identified in the cohort. Of this total, 67,416 peptides contained at least a single lysine residue, and we identified light-heavy (K0-K6) labelled peptide pairs in 41.25% of them, enabling the quantification of label incorporation in 27,752 peptides (Table S1). Overall, these data gave rise to 10,973 (± 392 sd) identified proteins and K6 incorporation was quantified in 2,685 (±190 sd) proteins per tissue sample. The change in turnover was calculated using proteins detected in at least two mice from each cohort, giving a screen that measured 1392 protein turnover changes in the *TgCRND8* hippocampus with high-quality MS data. A few proteins were absent from the screen because they contain very few lysine-containing peptides. One example of these proteins, ApoE, was of particular interest due to its association with AD. Therefore, to quantify these extremely scarce ApoE peptides, we enriched samples by immuno-affinity purification before MS analysis (Figure S1). In principle therefore, this *in vivo* approach can detect changes in protein turnover of any lysine-containing protein in any tissue.

Incorporation of K6 ranged from 1.4 to 70.9%, indicative of a large dynamic range of protein turnover. Remarkably, although there is minimal β-amyloid pathology at this age (Figure 1B and Table S1), an overall 6.1% slowdown in the global average protein turnover (GAPT) was measured in *TgCRND8* hippocampus (*P* = 0.0037 n = 1392, Figure 1C). In contrast, serum (*P* = 0.128, n = 118) and liver (*P* = 0.17, n = 590) protein turnover did not change significantly (Figure 1D-E), indicating that the decrease in protein turnover is specific to the pathologically affected forebrain tissue. As a further control to account for amino acid recycling rates, the precursor K6 concentration was determined using peptides containing more than one lysine in disease and control samples (Doherty et al., 2005; Hellerstein and Neese, 1992). No difference was detected (Table S2), indicating that changes in K6 incorporation directly measure changes in protein turnover.

If turnover changes are associated with β-amyloidosis, then as pathology progresses one would expect protein turnover changes to reflect the increasing load of β-amyloid pathology. To test this possibility, we repeated our turnover measurement at P285, by which age *TgCRND8* have pervasive amyloid deposits throughout the forebrain (Figure 2A, *top*, and Table S1). Surprisingly, examination of the protein turnover in the older *TgCRND8* model did not extend the slowdown that was seen at 113 days of age. Instead, at 285 days of age, an 18% increase in GAPT hippocampal protein turnover (*P* < 0.0001, n = 847, Figure 2A,B), whereas in serum protein GAPT was unchanged, indicating the change in proteome flux was restricted to the locus of β-amyloidosis. Overall, these proteome turnover data indicate discordance in proteome kinetics between early and late stages of pathology in the *TgCRND8* model of AD.

**Figure 2.**
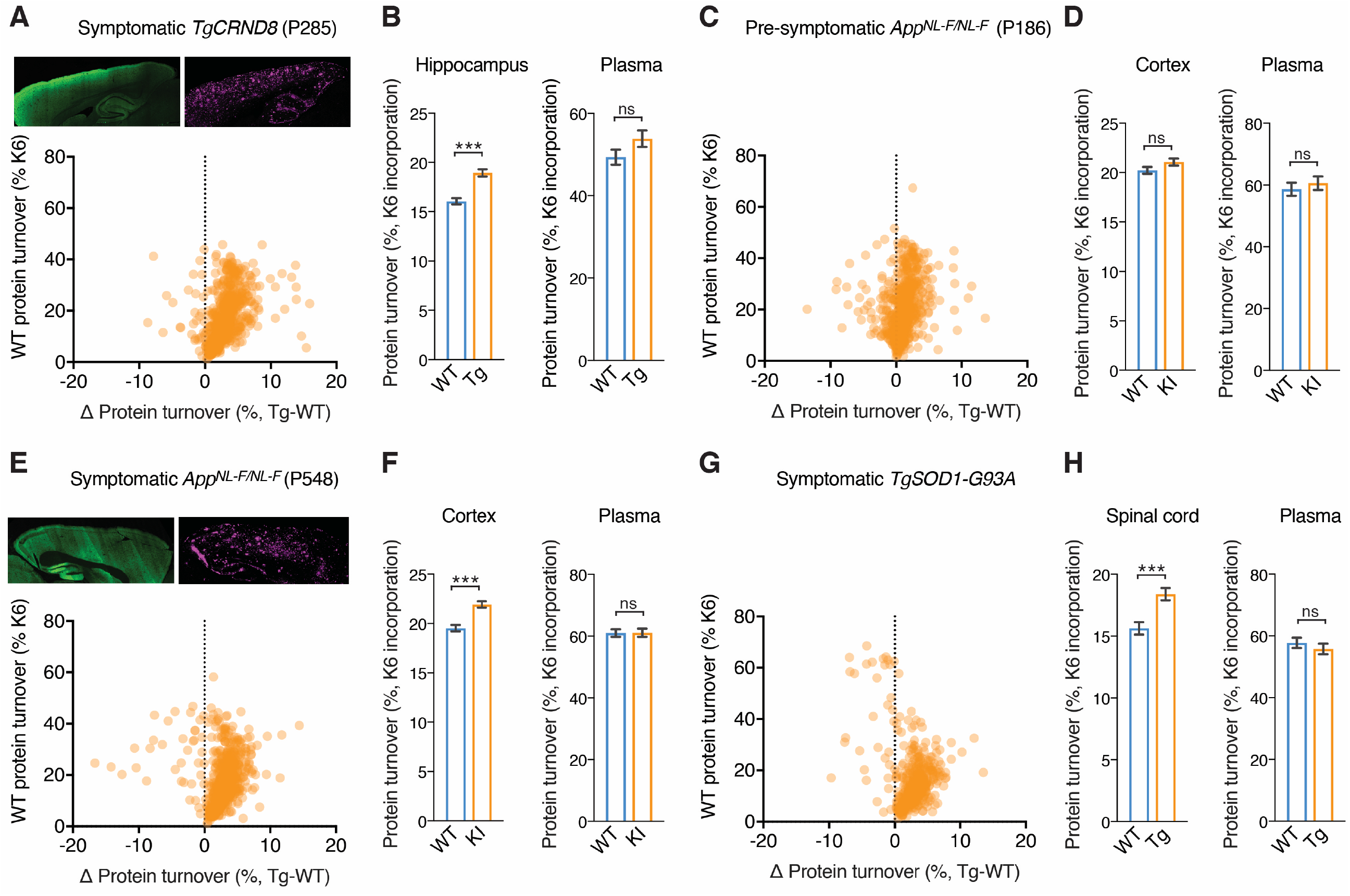
Differences in protein turnover in transgenic and knock-in models of AD, and a transgenic model of ALS. **A, *Top*,** Immunohistochemical detection of synaptic marker, left, Psd95 and right, β-amyloid pathology in sagittal sections of symptomatic *TgCRND8* (P285) mouse brain. ***Bottom*,** Scatter plot showing protein turnover changes in the hippocampus of symptomatic (P285) *TgCRND8* mice (see Table S1). 847 proteins were quantified, and the difference in turnover characteristics between proteins in diseased and healthy animals were plotted as described in Figure 1B. **B, *Left*** Bar chart showing the average hippocampal protein turnover of 847 proteins in *TgCRND8* and WT mice. An overall 18.0% increase in protein turnover was detected (*P* < 0.0001). Error bars indicate SEM. *** *P* < 0.0001. ***Right*** Bar chart showing average plasma protein turnover in *TgCRND8* and WT mice. No significant difference was detected (*P* = 0.102, n = 74). Error bars indicate SEM. ns, not significant. **C**, Scatter plot showing protein turnover changes in the cortex of pre-symptomatic (P186) *App^NL-F/NL-F^* mice (see Table S1). 721 proteins were quantified, and the difference in turnover characteristics between proteins in diseased and healthy animals were plotted as described in Figure 1B. **D, *Left*** Bar chart showing the average cortex protein turnover of 721 proteins in *App^NL-F/NL-F^* and WT mice. No change in protein turnover was detected (*P* < 0.0735). Error bars indicate SEM. ns, not significant. ***Right*** Bar chart showing average plasma protein turnover in *App^NL-F/NL-F^* and WT mice. No significant difference was detected (*P* = 0.534, n = 48). Error bars indicate SEM. ns, not significant. **E, *Top***, Immunohistochemical detection of synaptic marker, left, Psd95 and right, β-amyloid pathology in sagittal sections of symptomatic *App^NL-F/NL-F^* (P500) mouse brain. ***Bottom*,** Scatter plot showing protein turnover changes in the cortex of symptomatic (P548) *App^NL-F/NL-F^* mice (see Table S1). 721 proteins were quantified, and the difference in turnover characteristics between proteins in diseased and healthy animals were plotted as described in Figure 1B. **F, *Left*** Bar chart showing the average cortex protein turnover of 847 proteins in *App^NL-F/NL-F^* and WT mice. An overall 15.7% increase in protein turnover was detected (*P* < 0.0001). Error bars indicate SEM. *** *P* < 0.0001. ***Right*** Bar chart showing average plasma protein turnover in *App^NL-F/NL-F^* and WT mice. No significant difference was detected (*P* = 0.961, n = 95). Error bars indicate SEM. ns, not significant. **G,** Scatter plot showing protein turnover changes in the spinal cord of acutely symptomatic (P120) *TgSOD1-G93A* mice (see Table S1). 496 proteins were quantified, and the difference in turnover characteristics between proteins in diseased and healthy animals were plotted as described in Figure 1B. **H, *Left*** Bar chart showing the average protein turnover of 496 proteins in the spinal cord of *TgSOD1-G93A* and WT mice. An overall 17.6% increase in protein turnover was detected (*P* < 0.0001). Error bars indicate SEM. *** *P* < 0.0001. ***Right*** Bar chart showing average plasma protein turnover in *TgSOD1-G93A* and WT mice. No significant difference was detected (*P* = 0.417, n = 89). Error bars indicate SEM. ns, not significant.

### Proteome turnover in the *App^NL/F^* knockin mouse model of AD

In progressive diseases, identifying protein turnover changes that precede pathology could indicate upstream molecular pathways involved in the disease. However, in transgenic models of disease, including *TgCRND8*, one cannot distinguish between *bone fide* pathological mechanisms and effects that result from ectopic over-expression of the APP precursor. Therefore to test in the absence of over-expression, we used a knockin mouse model of familial AD, *App^NL-F/NL-F^*, which lacks these potential artefacts (Saito et al., 2014). Protein turnover was measured at P186 and P548, which are time-points before and after widespread β-amyloid pathology, respectively.

In pre-symptomatic P186 *App^NL-F/NL-F^* forebrain, no significant change in GAPT was detected (Figure 2C,D). However, in P548 *App^NL-F/NL-F^* mice with advanced β-amyloidosis, forebrain GAPT increased by 15.7% (*P* < 0.0001, n = 721), with 53 proteins being made or degraded faster (Figure 2E and Table S1). No significant change was detected in serum proteins (Figure 2F). Thus, an increase in protein flux is associated with increasing pathology in the *App^NL-F/NL-F^* knockin mouse model of β-amyloidosis.

### Protein turnover in tissue undergoing cell death

Late stages of neurodegenerative diseases are characterized by widespread neuronal death. To test for proteome turnover changes in tissues undergoing neuronal death we K6-labelled 3-month-old *TgSOD1-G93A* mice (Gurney et al., 1994), a mouse model of familial amyotrophic lateral sclerosis (ALS). At this age, *TgSOD1-G93A* mice displayed rear gait phenotypes (Gurney et al., 1994), indicating spinal cord pathology and extensive neurodegeneration. In *TgSOD1-G93A* spinal cord GAPT increased 17.7% in the diseased mice compared to control (*P* = 0.0001, n = 496, Figure 2G,H and Table S1). In contrast, plasma protein GAPT was unchanged (*P* = 0.417, n = 89). Overall, in all models at late stages a marked increase in overall protein flux was detected. These data could give insight into the particular pathways. However, this raises the question, whether or not increased turnover is coupled to the gain or loss in the expression levels of proteins, or if the increased turnover corresponds to the futile cycles of increased repair.

### Dynaplot: a comprehensive map of proteome dynamics

The kinetics of protein turnover drives the expression level of all proteins (Ohsumi, 2006). Therefore, the simultaneous measurement of turnover and expression level of each protein can give a comprehensive description of proteome dynamics and mechanistic insight. Having established a method for screening changes in protein turnover, we next combined turnover measurements with expression level measurements by exploiting recent improvements of label-free quantification (MaxLFQ) (Cox et al., 2014). In each mouse model of disease dataset, an average of 4357 proteins (± 654 sd) were quantified by label-free quantification (Table S3). Proteins were quantified in all mice in 98.0% (±2.0%) of the proteins that were used for turnover analysis, giving excellent reproducibility.

Plotting turnover versus steady state expression level changes (hereon referred to as a dynaplot) is potentially a powerful tool because the coordinate space of these measurements infers the mechanism of change, as depicted in Figure 3A. In principle, changes in turnover can be regulated by either the rate of synthesis or degradation. Therefore, six scenarios arise: 1) Increased steady state levels driven by an increase in protein turnover, indicates the net synthesis rate has increased. 2) Increased steady state levels can also be driven by a net decrease in turnover, which reflects a decrease in the rate of protein degradation. Similarly, two distinct mechanisms for decreasing the steady state level of proteins can be directly inferred: 3) Decreased steady state levels driven by a decrease protein turnover, which is the result of a net decrease in protein synthesis, and 4) decreased steady state levels driven by an net increase in protein turnover, which is driven by a net increase in degradation. Finally, proteins can occupy coordinate space on the dynaplot in which 5) increases or 6) decreases in protein turnover are uncoupled from steady state changes. These futile cycles reflect an increase and decrease in the rate of protein repair, respectively.

**Figure 3.**
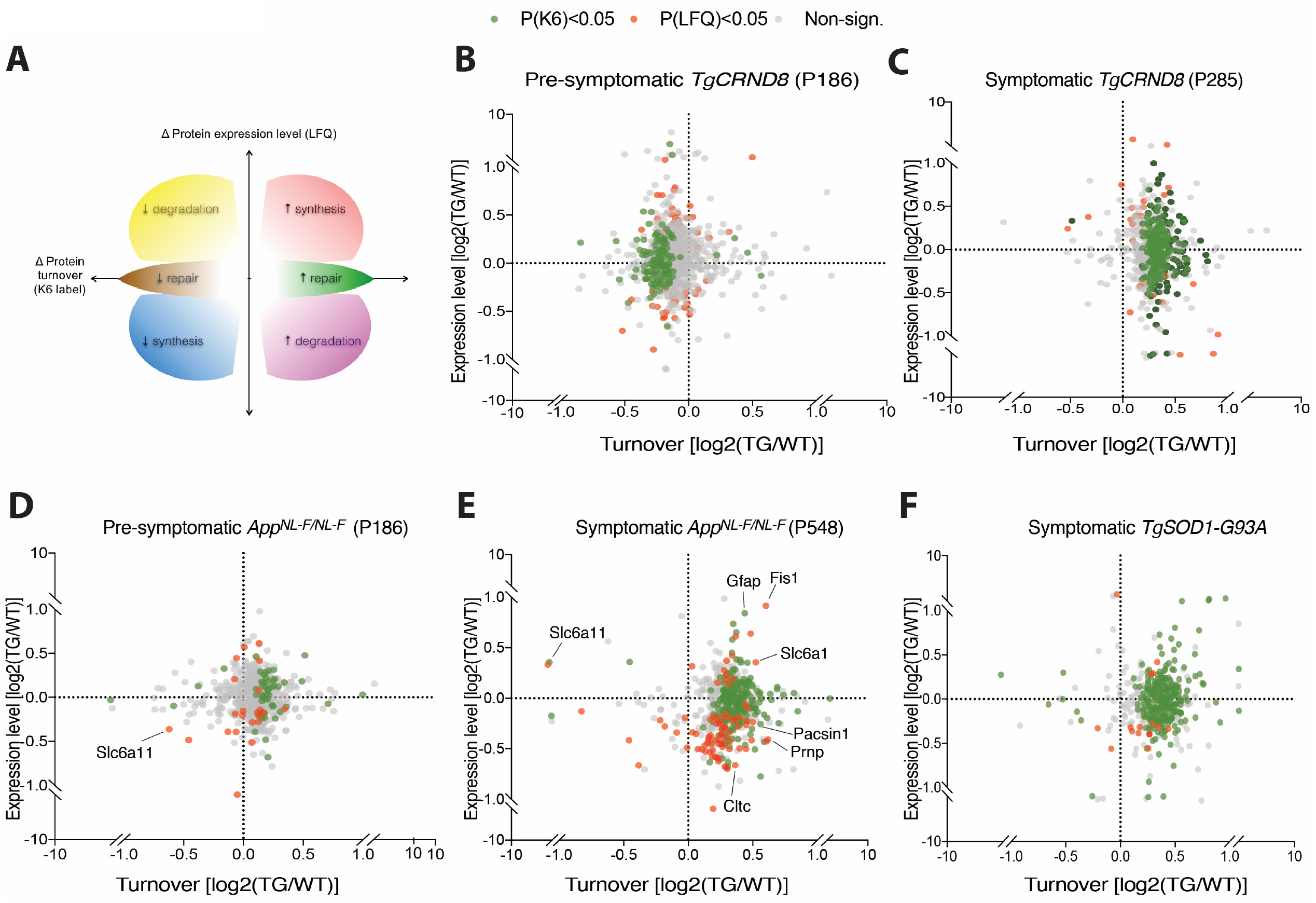
Multidimensional measurement of proteome dynamics in live mice. **A,** Schematic of a dynaplot showing the change in protein turnover (x-axis, K6 label) plotted against the change in steady state expression level (y-axis, LFQ). The coordinate space of the plot reflect dynamics of a protein that can be attributed to a net increase (red) or decrease (blue) in the rate of synthesis; net increase (magenta) or decrease (yellow) in degradation. Change in turnover that does not result in a change in steady state expression indicate change in flux of increasing (green) or decreasing repair (brown). **B,** Dynaplot of hippocampal proteins in pre-symptomatic (P113) *TgCRND8* mice, as compared to healthy, matched control mice. Proteins that were significantly different in turnover (*P* < 0.01) are highlighted in green, while those that were significantly different in steady state amount (*P* < 0.01) are highlighted in red. **C,** Dynaplot of hippocampal proteins in acutely symptomatic (P285) *TgCRND8* mice, as compared to healthy, matched control mice. 4. **D,** Dynaplot of cortex proteins in pre-symptomatic (P186) *App^NL-F/NL-F^* mice, as compared to healthy, matched control mice. **E,** Dynaplot of cortex proteins in symptomatic (P548) *App^NL-F/NL-F^* mice, as compared to healthy, matched control mice. **F,** Dynaplot of hippocampal proteins in acutely symptomatic (P120) *TgSOD1-G93A* mice, as compared to healthy, matched control mice. See Table S 4.

A dynaplot showing the change in flux versus the expression level of each protein is depicted in Figure3. In each disease mouse model imbalances in turnover resulted in changes in expression level of a subset of the proteome (Figure 3B-F). Comparing 6-month and 18-month-old *App^NL-F/NL-F^* showed a 5-fold increase in the number of significantly changed proteins (Figure 3D,E and Table S3). Thus, increased pathology correlated with increased imbalances in the proteome. However, different proteins were differentially affected, suggesting different pathways are impacted at early and late stages of pathology (Table S4). Analysis of the symptomatic *App^NL-F/NL-F^* dataset using the KEGG showed that significantly changed proteins converged on several pathways that appear to be prevalent in presynaptic functions, including synaptic vesicle recycling and mitochondria (Figure 4). Thus, multidimensional proteome dynamics have identified specific proteins and pathways dysregulated as a consequence of disease (Table S3).

**Figure 4.**
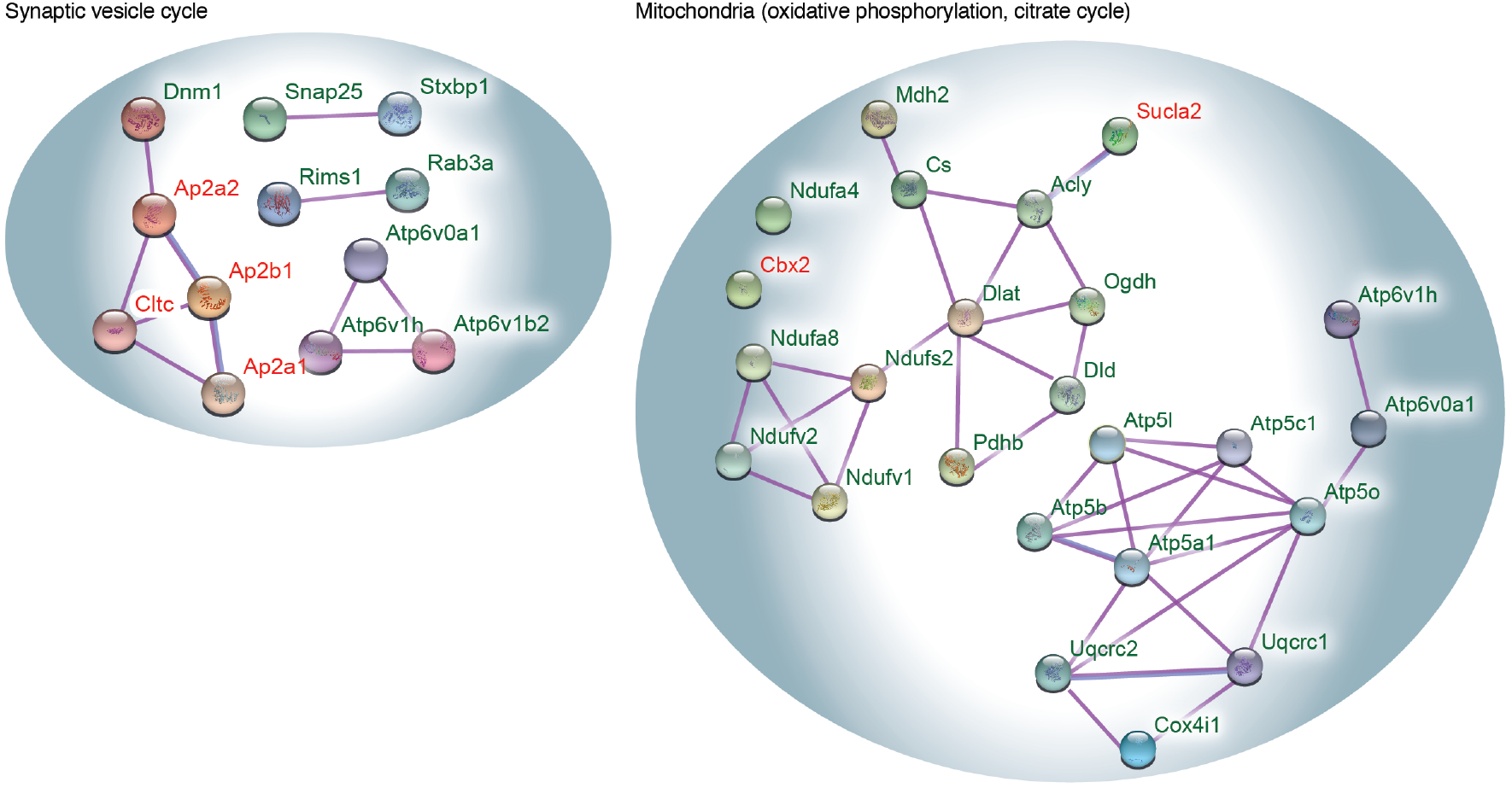
Proteins with perturbed dynamics converge on presynaptic functions in mouse model of Alzheimer’s disease. Gene ontology enrichment using KEGG database (see methods) identified left, synaptic vesicle recycling, and right, mitochondrial pathways enriched with the proteins whose turnover (green) or steady state amount (red) have significantly changed in symptomatic (P548) *App^NL-F/NL-F^* cortex. Purple and blue edges, indicate empirically determined protein-protein interactions, and protein homology, respectively.

### Proteome dynamics of ageing and autophagic flux

Ageing is the greatest known risk factor for developing neurodegenerative disease, including AD (Blennow et al., 2006). It is commonly held that protein turnover slows with age (Dwyer et al., 1980; Makrides, 1983). Therefore, we measured the change in proteome dynamics associated with ageing in wildtype mice varying in age from 3 to 17 months (Figure 5A and Table S5). Strikingly, 98% of the proteins measured showed decreasing K6 incorporation with age (between 113 and 503 days of age, *P* = 6.2×10^−16^, n = 360). Thus, the global average protein turnover decreased significantly with age (Figure 5A). Since the K6 label is delivered by diet, as a control we measured average food consumption, which indicated no decline in food consumption with age (Figure 5B). The average levels of protein abundance did not change significantly (Figure 5C and Table S5), consistent with earlier reports (Walther and Mann, 2011). Overall, these ageing data could provide a rich resource for exploring molecular mechanisms associated with ageing.

**Figure 5.**
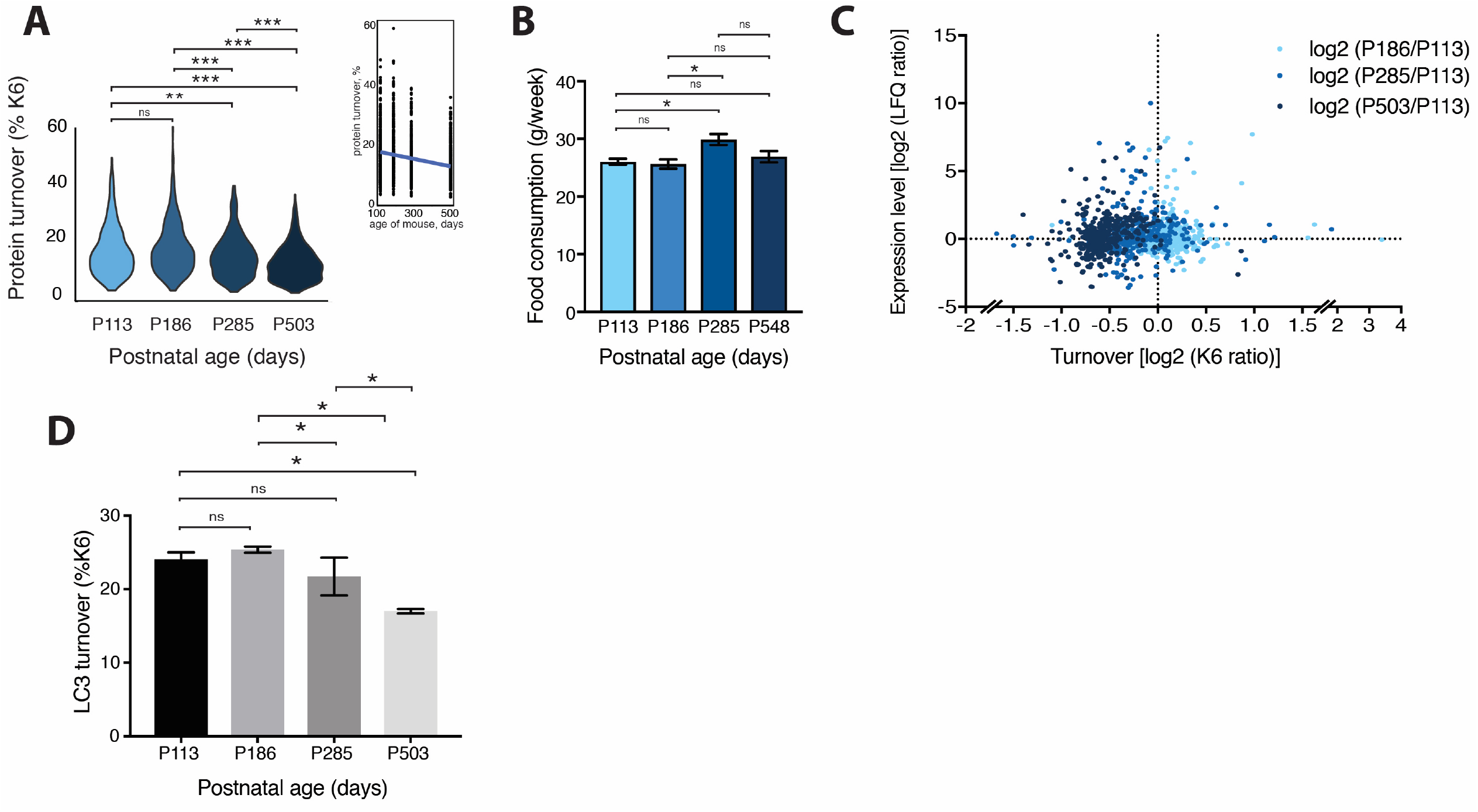
Proteome dynamics associated with ageing in healthy mice. **A,** Average protein turnover in healthy mouse cortex at various ages. Total number of proteins used to compare turnover across all ages = 360. P values for the comparisons (descending from top) = 1.063E-06, 4.557E-14, 1.220E-05, 6.799E-14, 0.003, 0.175. **B,** Average food consumption of mice at different ages: P113 (n=4), P186 (n=2), P285 (n=4), P548 (n=11). No significant change in appetite associated with ageing was detected. P values for the comparisons (descending from top) = 0.127, 0.675, 0.047, 0.694, 0.011, 0.692. **C,** Dynaplot of mouse cortex proteins depicting the change in proteome dynamics in healthy control mice from P113 to P186 (sky blue), to P285 (blue), and to P548 (dark blue). These data are expressed ratiometrically to allow simpler visualisation between different ages of mice. **D,** Bar graph showing apparent decrease in turnover of LC3, marker of autophagy, as mice age. Turnover was estimated by percentage heavy lysine incorporation (P6) in mice at postnatal days 113, 186, 285 and 503. * *P* < 0.05. n.s., non-significant.

Next, we explored our proteome dynamics dataset to identify potential mechanisms that could explain the ageing-associated global decrease in proteome turnover, focusing on mechanisms that could drive proteome wide changes. One such candidate is autophagy, the dynamics of which are challenging to follow (Loos et al., 2014; Schmidt et al., 2009). A prominent marker of autophagy is LC3, and this showed a significant decrease in K6 incorporation that was tightly associated with ageing (Figure 5D). However, the steady state levels of LC3 fluctuated but did not correlate with increasing age (Table S5). Thus, autophagic flux decreases with age and provides a mechanism that in part explains the global decrease in proteome turnover.

## DISCUSSION

Using a novel combination of live mouse labelling and proteomic profiling, we have developed a method for simultaneously measuring the flux of proteome wide changes in turnover and steady-state expression levels. By combining both measurements, this approach distinguishes changes that are mediated protein synthesis versus degradation and enables the direct estimate of changes in protein flux. We tested this method in multiple mouse models of disease, which revealed global effects as well as identifying individual pathways associated with pathology. Next, we applied the method to ageing in wildtype mice, and observed a decline in protein turnover that could be partially explained by a decline *in vivo* autophagic flux.

This methodological framework should be especially useful for identifying proteins and molecular pathways in other live animal test settings, including disease models, learning and behaviour (Shen et al., 2014). Establishing the cause of an imbalance in turnover could be essential for understanding diseases, and is likely to highlight new targets for pharmaceutical intervention. Indeed, there is a growing body of evidence that correcting imbalances in proteome dynamics can slow the onset of disease symptoms (Das et al., 2015; Moreno et al., 2012).

There is also increased appreciation that better mouse models are needed to identify targets for therapeutic intervention in neurodegenerative diseases, including AD (Drummond and Wisniewski, 2017). The comprehensive proteome dynamics provided insights that enable the direct comparison of multiple different mouse models (Savas et al., 2017). Comparing the over-expressing hAPP transgenic model (*TgCRND8*) at early and late stages of the disease indicated large yet discordant effects on protein turnover. Since changes did not correlate with increasing pathology it is difficult to distinguish molecular mechanisms altered by over-expression of the transgene from changes associated with β-amyloidosis in this mouse line. In contrast, turnover changes identified in models that do not rely on ectopic over-expression, (*App^NL-F/NL-F^* knockin model) were correlated with increased pathology.

In all models tested at symptomatic stages, global average protein turnover increased suggesting a disease-associated proteome-wide state of repair. Increased proteomic flux drove expression level imbalances caused by increased synthesis of one subset of the proteome and increased degradation of another (Figure 3 and Table S4). This is consistent with transcriptomic data from AD postmortem samples that suggested increased autophagy-mediated turnover (Lipinski et al., 2010). It is likely transcriptional programmes are involved in regulating the increase in proteome flux (Matarin et al., 2015). Also, an increased flux of Aβ has been detected in familial AD patients (Potter et al., 2013).

In contrast to disease, it is intriguing that turnover declines as wildtype mice age, whereas steady state levels of the proteome in mice appear to show no global change associated with age (Walther and Mann, 2011). This is consistent with similar reports of proteome turnover decline in invertebrates (Walther et al., 2015), albeit expression levels in ageing invertebrates appear to change (Narayan et al., 2016; Walther et al., 2015). Our turnover measurements of LC3 enabled an estimation of autophagic flux, which also declined with ageing, suggesting the global turnover decrease associated with ageing could, at least in part, be mediated by slowdown in autophagy.

Overall, it is striking that while ageing is associated with a decrease in flux, all three neurodegenerative disease models caused an increase in flux. Thus, these mouse models suggest neurodegenerative diseases are not an acceleration of ageing, but rather represent a state of proteome imbalance and increased repair. As improved models of neurodegenerative disease are developed, applying comprehensive proteome dynamics is expected to give important phenotypic, molecular, and mechanistic insight.

### STAR methods

KEY RESOURCES TABLE

KEY RESOURCES TABLE
| REAGENT or RESOURCE | SOURCE | IDENTIFIER |
| --- | --- | --- |
| **Antibodies** |  |  |
| Mouse monoclonal (4G8) anti-β-Amyloid (17-24) | BioLegend | Cat# 800701<br>AB_2313891 |
| Mouse monoclonal (6E10) anti-β-Amyloid (1-16) | BioLegend | Cat# 803015<br>AB_10102028 |
| Rabbit polyclonal anti-PSD95 | Abcam | Cat# ab18258<br>AB_444362 |
| Rabbit monoclonal (16H22L18) anti-ApoE | Thermo | Cat# 701241<br>AB_2532438 |
| Mouse monoclonal (D6E10) anti-ApoE | Abcam | Cat# ab1906<br>AB_302668 |
| **Deposited Data** |  |  |
| Raw and database-matched mass spec data | This paper | PRIDE: PXD010671 |
| **Experimental Models: Organisms/Strains** |  |  |
| *TgCRND8* | David Westaway | PMID: 11279122 |
| *App<sup>NL-F</sup>* | Takaomi Saido | PMID: 24728269 |
| *TgSOD1-G93A* | Jackson Labs | PMID: 8209258 |
| **Software and Algorithms** |  |  |
| MaxQuant | Cox and Mann, 2008 | http://www.coxdocs.org/doku.php?id=maxquant:common:download_and_installation |
| Perseus | Tyanova *et al*, 2016 | http://www.coxdocs.org/doku.php?id=perseus:common:download_and_installation |
| Prism | Commercial software | https://www.graphpad.com/scientific-software/prism/ |
| R | Open source | https://www.r-project.org/ |

## CONTACT FOR REAGENT AND RESOURCE SHARING

Further information and requests for resources and reagents should be directed to and will be fulfilled by the lead contact René A. W. Frank

## EXPERIMENTAL MODEL AND SUBJECT DETAILS

### Mice

Animals were treated in accordance with UK Animal Scientific Procedures Act (1986) and NIH guidelines. Several mouse models of neurodegeneration were used in this work. Two models of β-amyloidosis were used, the TgCRND8 transgenic mouse and the *App^NL-F/NL-F^* knock-in mouse (Chishti et al., 2001; Saito et al., 2014). The *SOD1-G93A* mice modelled ALS (Gurney et al., 1994). Genotyping was performed on every mouse by the LMB in-house facility. Age-, sex- and genetic background matched mice were used in test and control groups.

## METHOD DETAILS

### Metabolic labelling of live mice

Mouse proteins were globally labelled ^13^C heavy lysine (Lysine-6; K6) by feeding with mice Lys-6 food (Silantes, Germany) for 6-8 days. All transgenic models of disease were fed K6 food for 6 days, as were the control mice used in the ageing analysis. To counteract the decline in protein turnover that is observed in age (Figure 5), the 18-month-old *App^NL-F/NL-F^* and control of mice were fed for eight days. Mice were kept in cages separated by genotype, and labelled in groups of six: three control and three experimental mice. The mass of each mouse and food consumed were recorded throughout the experiments and the primary control for assessing the level of heavy label incorporation was the measurement of plasma protein turnover.

At the end of the labelling period, the mice were culled and all major organs were collected and immediately frozen in liquid nitrogen, including blood plasma and cerebrospinal fluid. The brain was dissected into eight separate regions, and each area was frozen individually and immediately (olfactory bulb, caudate putamen, hippocampus, cortex, colliculus, cerebellum, thalamus and hindbrain).

### Mouse tissue fractionation

Mice were culled by cervical dislocation or by overdose of pentobarbital. Organs were immediately dissected on ice, including the separation of brain areas and harvesting of cerebrospinal fluid, and all tissue and humours were snap frozen in liquid nitrogen. Tissue was homogenised manually in H buffer (320 mM sucrose, 2 mM HEPES at pH 7.3 with protease inhibitors). Volumes of H buffer were scaled to mass of tissue (232 mg tissue = 5 ml H buffer). Nuclei were pelleted by centrifugation at 1k x g, and this pellet was homogenised for a second time in H buffer and pelleted as above. Membranes were pelleted from the supernatant at 21k x g and all fractions were divided into small portions and flash-frozen in liquid nitrogen. This procedure was applied to hippocampus, cortex and spinal cord. Plasma proteins were diluted 1:40 with PBS and added to LDS sample buffer (Thermo) directly. For extraction, tissue fraction pellets were resuspended in H buffer, and extraction/precipitation buffer added to the suspension as appropriate. Extraction/precipitation buffer consisted of 25 mM Tris, pH 8, 50 mM NaCl, 2 mM TCEP, protease inhibitors, benzonase (Novagen) and detergent - proteins from the membranes were solubilised in deoxycholate or Triton X-100 (0.8% w/v or 1% v/v final, respectively) and proteins from the nuclear fraction were precipitated in Triton X-100 or GDN (1% final). Solubilised material was cleared by ultracentrifugation at 120k x g for 40 mins, 8C and precipitated material was pelleted by centrifugation at 21k x g for 25 mins, 8C. Solubilised or precipitated material was prepared for SDS-PAGE by addition of LDS sample buffer and cysteines were alkylated with 10 mM iodoacetamide prior to electrophoresis through 1 mm thick 4-12% Bis-Tris acylamide gels (Thermo). Proteins were stained with colloidal Coomassie blue, and 16 sections, each 4 x 4 mm, were cut from each sample lane and diced into 1 mm cubes separately. These polyacrylamide cubes containing the fractionated proteins were prepared for mass spectrometric analysis using the Janus liquid handling system (PerkinElmer, UK). Briefly, the excised protein gel pieces were placed in a well of a 96-well microtitre plate and destained with 50% v/v acetonitrile and 50 mM ammonium bicarbonate, reduced with 10 mM DTT, and post-alkylated with 55 mM iodoacetamide. After alkylation, proteins were digested with 6 ng/μL trypsin (Promega, UK) overnight at 37 °C. The resulting peptides were extracted in 2% v/v formic acid, 2% v/v acetonitrile. In some cases, the peptides extracted from 16 gel sections were combined into 4 samples for LC-MS. The conditions of detergent fractionations and digestion with trypsin were chosen from several rounds of optimisation experiments to maximise peptide coverage and quantification.

### Mass spectrometry and data analysis

The protein digest was analysed by nano-scale capillary LC-MS/MS using an Ultimate U3000 HPLC (ThermoScientific Dionex, San Jose, USA) to deliver a flow of approximately 300 nL/min. A C18 Acclaim PepMap100 5 *μ*m, 100 *μ*m x 20 mm nanoViper (ThermoScientific Dionex, San Jose, USA), trapped the peptides prior to separation on a C18 Acclaim PepMap100 3 *μ*m, 75 *μ*m x 250 mm nanoViper (ThermoScientific Dionex, San Jose, USA). Peptides were eluted with a 120 minute gradient of acetonitrile (2% to 50%). The analytical column outlet was directly interfaced, via a nano-flow electrospray ionisation source, with a hybrid quadrupole orbitrap mass spectrometer (Q-Exactive Plus Orbitrap, ThermoScientific, San Jose, USA). Data dependent analysis was carried out, using a resolution of 30,000 Da for the full MS spectrum, followed by ten MS/MS spectra. MS spectra were collected over a m/z range of 300–2000. MS/MS scans were collected using a threshold energy of 27 for higher energy collisional dissociation (HCD). Each tryptic peptide containing lysine-6 produced a peptide ion pair differing by 6.02 Da (divided by charge state).

For SILAM analysis of protein turnover, peptide pairs were located with MaxQuant 1.5.0 and identified with Andromeda using a reviewed version of the mouse Uniprot database. Each detergent fraction was analysed with MaxQuant individually, and quantified proteins were forwarded for analysis if they were found in at least two of three biological replicates in both control and experimental animals. A final non-redundant merged dataset was generated excluding quantifications of the same protein from different detergent fractions; keeping the protein measurement with the greatest difference in Lysine-6 incorporation between the control and experimental animals. Two-tailed Student’s T tests (p = 0.05) were performed on the ratios of incorporation in control and experimental animals. As detailed in figure 1a, the SILAM ratio is a direct readout of protein turnover.

For Label-Free Quantification of the same datasets, the maxLFQ functionality of MaxQuant 1.5.0 was used. However, proteins in each detergent fraction were forwarded for subsequent analysis if they were found in every single mouse of that fraction (six of six). The protein quantification data were integrated by averaging the quantities from each high-quality detergent fraction together on a mouse-by-mouse basis. Two-tailed Student’s T tests (p = 0.05) were performed on the levels of protein in control and experimental animals.

Importantly, it was essential to observe the peptide pairs at least twice, by using a double count requirement for quantification in MaxQuant. We found experimentally that if the pairs of peptides were measured only in only a single scan, peptide ratios were erroneously found in over 2% of identified proteins in an unlabelled sample (57 ratios in 2733 identifications, data not shown).

### Immunohistochemistry

Mice were euthanized by cervical dislocation or by overdose of pentobarbital when they were transcardially perfused with PBS. Brains were divided along the midline and half was submerged in OCT (optimal cryotomy) solution in a cut-away plastic mould – the other half was kept for biochemical analysis. OCT-submerged brains were frozen by submersion of the mould into a beaker of isopentane that was subsequently chilled in liquid nitrogen. The frozen brain sections were cut at a thickness of 14 um using a cryostat (Leica) and post-fixed using freezing methanol. Sections were blocked using 3% BSA or 10% goat serum in PBS with 0.2% Triton X-100, and probed with primary antibodies [Mouse anti-Abeta (6E10; Covance, 803001) and Rabbit anti-PSD95 (Abcam, ab18258] overnight at 4 C. Secondary antibodies were conjugated to Alexafluor 488 or 647 (Thermo) and were applied to the samples for two hours before mounting the slide using ProLong Antifade mountant with DAPI (Thermo). Images were acquired using a Zeiss 780 confocal microscope then viewed and analysed in Fiji. All antibody combinations were validated by controls with individually absent primary antibodies.

### Immuno-affinity protein purification

Frozen membrane fractions were resuspended in H buffer and solubilised in DOC extraction buffer (as above) for 1 hour. The extract was cleared by ultracentrifugation (120k x g, 40 mins, 8C). Antibodies were added to the cleared extract and left to bind overnight at 4C. The antibodies and adsorbed proteins were reclaimed by the addition of protein G Dynabeads (Sigma) for 40 mins the following day. After washing twice with a solution containing 25 mM Tris, pH 8 and 50 mM NaCl, the antibodies and the adsorbed proteins were eluted with SDS.

### Functional clustering analysis

Proteins that changed their turnover dynamics were functionally clustered by the DAVID online tool (https://david.abcc.ncifcrf.gov/home.jsp), using KEGG pathways. Functional annotation charts were exported and visualized using String (Szklarczyk et al., 2017) to depict experimentally determined protein-protein interactions.

## QUANTIFICATION AND STATISTICAL ANALYSIS

### Statistics

Data were analyzed with PRISM 5 (GraphPad). Data are presented as mean and error bars represent SEM. All experiments were performed in at least three independent biological replicates. Details of statistical tests used and p values are presented in the figure legends.

*p ≤ 0.05, **p ≤ 0.01, ***p ≤ 0.001, ****p ≤ 0.0001, ns: non-significant.

## DATA AND SOFTWARE AVAILABILITY

### Proteomic data deposition

All data were deposited in the PRIDE database, project accession: PXD010671.

## Acknowledgements

We thank the staff of the MRC Laboratory of Molecular Biology’s Animal Facility. We would also like to thank Nick Barry, Mathias Pasche, and Jonathan Howe for light microscopy support. Jake Grimmet and Toby for computational infrastructure support. Sebastian Schmidt (Silantes) for technical support with SILAM labelling. We would also like to thank Michel Goedert, Harvey McMahon, Nushan Pasindu Gunawardana, Toke Hansen, Isabelle Lavenir, Jennifer McDonald and Ben Falcon (MRC LMB) for insightful discussions. The research leading to these results received core funding from the Medical Research Council and a pump priming grant from Alzheimer’s Research UK.

